# Integrative functional analysis uncovers metabolic differences between *Candida* species

**DOI:** 10.1101/2021.05.24.445215

**Authors:** Neelu Begum, Sunjae Lee, Aize Pellon, Shervin Sadeghi Nasab, Jens Nieslen, Mathias Uhlen, David Moyes, Saeed Shoaie

## Abstract

*Candida* species are a dominant constituent of the human mycobiome and a better understanding of their metabolism from a fungal perspective can provide key insights into their ability to cause pathogenesis. Here, we have developed the BioFung database – a fungal specific tool for functional annotation using the KEGG database that provides an efficient method for annotation of protein-encoding gene. Analysis of carbohydrate-active enzyme (CAZymes) and BioFung, uncovered core and accessory features across *Candida* species demonstrating plasticity, adaptation to the environment and acquired features. Integerative functional analysis revealed that all *Candida species* can employ amino acid metabolism. However, metabolomics revealed that only a specific cluster of species (AGAu species - *C. albicans, C. glabrata and C. auris*) utilised amino acid metabolism. We identified critical metabolic pathways in the AGAu clusters with biomarkers and anti-fungal target potential in the CAZyme profile, polyamine, choline and fatty acid biosynthesis pathways. This study, combining genomic analysis, metabolomics and gene expression validation, highlights the metabolic diversity within AGAu species that underlies their remarkable ability to dominate the mycobiome and cause disease.

## Introduction

Fungal infections affect around 7.5 million people around the world every year. Within human fungal communities (mycobiome), with the notable exception of the skin, *Candida* species are the most common group^1–3^. These species are generally pathobionts, being the most common human fungal pathogens, despite also being commensal organisms^4^. *Candida* infections are becoming increasingly concerning, and the World Health Organisation (WHO) has recently emphasised international surveillance for diagnosis and management of fungal infection, particularly *C. albicans* infection^5–7^. Recently, a novel *Candida species, Candida auris* has been identified with significant mortality and morbidity, as well as a high degree of antifungal resistance^8,9^. There are over 200 *Candida* species currently identified, but only a handful of these are present in the human microbiota with the ability to cause infection and pathology. Most notable examples of these include *C. albicans, C. glabrata, C. dubliniensis, C. tropicalis* and *C. auris*^9–15^. *Candida* species, most notably *C. albicans* and *C. glabrata*, can give rise to a variety of superficial infections, including oral thrush and vulvovaginal candidiasis, but are also capable of causing a systemic infection with significant mortality^16–20^. As well as direct infections, fungi such as *Candida* species have also been associated with oncogenesis through complement activation, demonstrating potential effects of the interaction of fungal species with human host ^21^.

An essential virulence determinant of fungi is their metabolic plasticity^22^. Fungal are significant in their ability to utilise numerous different anabolic and catabolic sources in their metabolic processes, attributable to switching between carbon and nitrogen sources^23^. Nutritional availability, environmental factors, competition and pathogenic factors all influence this plasticity^24,25^. Investigations of *Candida* species-specific transcriptional regulators of glycolytic genes (e.g. Tye2 and Gal4) and enzymes of the glycolytic pathway (hexose catabolism), indicate these factors play an essential role in central carbon metabolism commonly applied during infection events^22,24,26^. Glycolytic metabolism can activate virulence factors that initiate hypha formation, activate fermentative pathways, repress gluconeogenesis, and the TCA cycle^27–30^. Alternatively, *C. albicans* can switch to gluconeogenesis and the glyoxylate cycle to confer full pathogenesis during systemic candidiasis^31–34^. Carbohydrate metabolism is coupled with changes of cell wall architecture, host immune response modulation, as well as adherence, biofilm formation, stress response and drug resistance^24,35–37^. If carbohydrate sources are limited, *Candida* species can use amino acids and lipids as supplementation for metabolic adaptation^22,38–40^. Amino acids produced by *C. albicans* have been shown to drive tissue damage by initiating stress responses and adjusting the surrounding environmental pH, helping induce of host invasion processes ^35,41–47^. Very little is known about the regulation, process and utilisation of amino acid metabolism in *Candida*^39^. However, *C. albicans* is known to use amino acids to replace carbon and other nitrogen sources^48^. *Candida*’s ability to convert arginine to urea allows the neutralisation of an acidic environment triggering the development of hyphae and biofilm formation ^32,49,50^. Notably, recent work has shown that *C. albicans* phagocyted by macrophages induces fatty acid β-oxidation and the glyoxylate pathway to induce hypha formation for escape. In a harsh environment that lacks even a nitrogen source, *Candida* can recycle and produce its own proteins and polyamines without host nitrate^51^. Thus, understanding of metabolism and functionality of *Candida* is instrumental in tackling infection and mortality prevalence^52^.

Here we developed BioFung – a database focused on functional information and interpretation of biological information. It is based on 128 fungal species using KEGG orthologs (KO). By applying BioFung to *Candida species*, we identified a distinct cluster of *C. albicans, C. glabrata* and *C. auris* referred to as AGAu species. This cluster has a high association with infection and mortality, relative to other *Candida* species^18,53–55^. We applied comparative analysis techniques based on gene, protein, and enzyme-substrate levels and identified metabolic pathways in *Candida* species, such as choline and polyamine pathways. Metabolomics along with experimental validation from gene expression confirmed the AGAu cluster difference. This study (1) provides a novel tool for functional annotation of fungal species, (2) highlights amino acid metabolism importance in AGAu species, and (3) identifies novel potential fungal biomarkers and anti-fungal targets in metabolic pathway.

## Results

### Development of BioFung database and functional annotation of *Candida* protein-encoding genes

To perform a global functional analysis of *Candida* species, we collected 49 publicly available genomes of different *Candida* strains covering 13 different species (Supplementary Table 1). We selected species based on their clinical importance and abundance within the human mycobiome^56–60^. All 49 *Candida* strains were isolated from different body sites from people in different geographical locations (Fig. 1a and Extended Fig. 1a). The quality of the genome of each strain was checked at the level of scaffold assembly and genome size (Extended Fig. 1b before reassessing the phylogenetic relationship of these 49 strains based on nucleotide sequence using average nucleotide identity (ANI) (Fig. 1b, Method). We observed strain-specific differences in phylogenetic lineages with 11 distinct branches, including branching of *C. auris, C. glabrata* and *C. albicans*, implying genetic diversity that could implicitly be interpreted into functional variances. To elucidate functional details for these strains, we built fungi-specific Hidden Markov Models (HMM) using fungal gene clusters, named BioFung database (Fig. 1c, Method)^61,62^. We analysed 524,288 fungal genes, from 128 fungal species, with a coverage of 4,822 KOs, and 4,430 fungal KO alignments to create BioFung (Supplementary Table 2-4). Comparison of the BioFung database with other eukaryote-specific HMM sources indicated that the BioFung has both higher coverage and specificity of KOs (Extended fig. 1c-e) ^61,62^.

**Figure 1.**
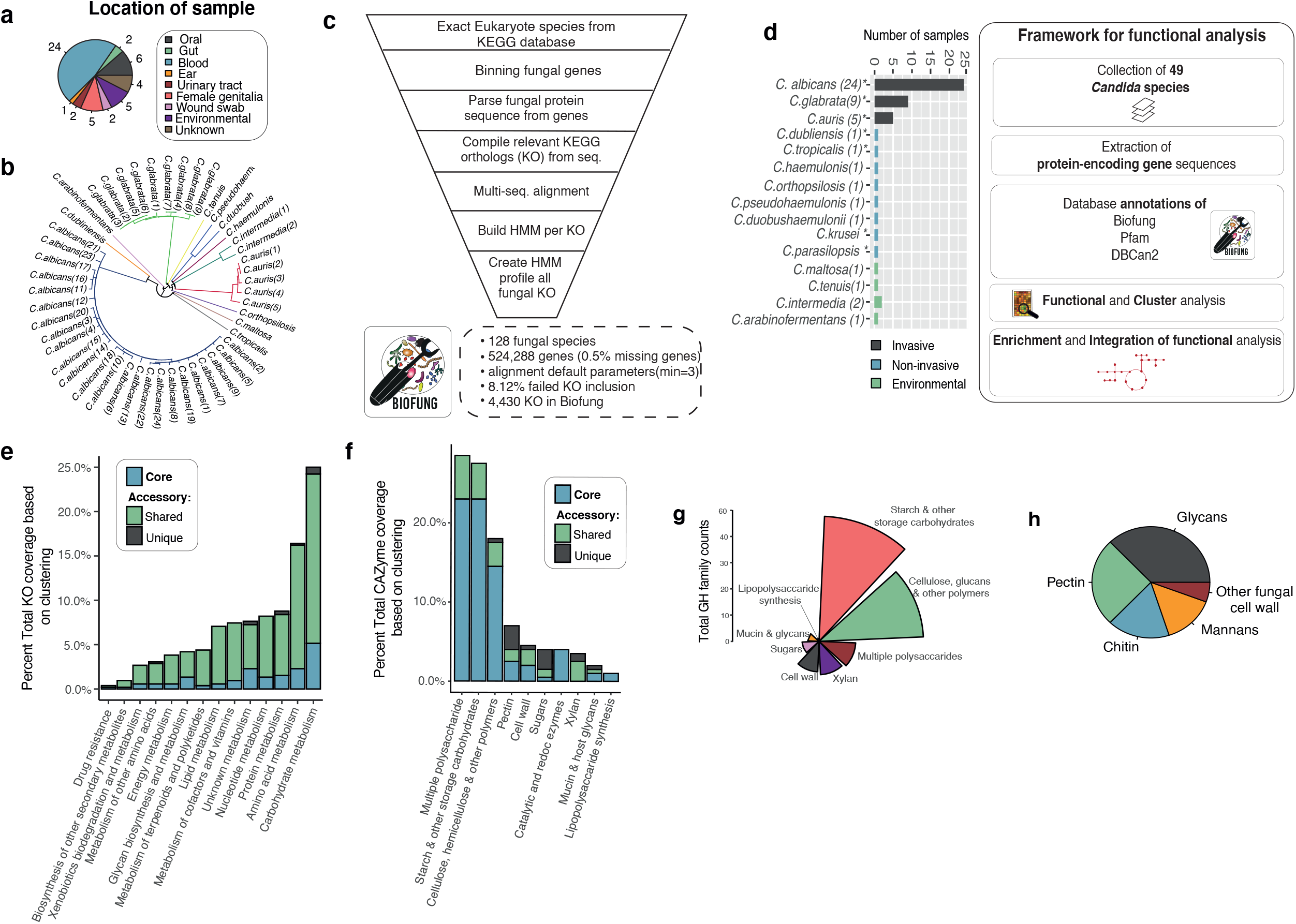
a, *Candida* strain characterisation. Coverage of *Candida* sample population per species available with the categorisation of species profiled. Numbers around the pie chart signify the number of strain representation in each location. (Supplementary Table 4 for more information about strains and extended figure 1a for the global representation of Candida strains). b, Genome-based phylogenetic tree. The phylogenetic tree was constructed based on average nucleotide identity (ANI) between all strains revealing evolutionary differences across strains (colour coordinated) and indicating distinct metabolic capabilities. See Extended Fig. 1d for quality of sequences. c, BioFung database creation workflow. Eukaryote annotation from KEGG database parsed to extract all fungal species. They were genes parsed, sequences extracted and reassembled to KO. The multi-sequence alignment was performed on each KO with all corresponding sequence available. HMM, profile built based on each KO and assembled to provide a more accurate annotation of fungal species for KO. d, Distribution of *Candida* species based on sample collection and the framework of protein-encoded genes analysis of *Candida* strains. Strains isolated from the various location providing relevant clinical association to host mycobiome and environmental species. * indicates clinical strains used for metabolomics. Functional analysis performed on 49 *Candida* species collected from public repositories. Protein sequences were annotated with Pfam, dbCAN2 and BioFung database for biological information. e, Core and accessory overview of the metabolic pathway across *Candida* strains. Shared genome feature refers to 6-48 species sharing the function and unique genome features is exhibited by less than 5 *Candida* species denoting accessory functions. f, Clustering of carbohydrate-active enzyme profile (CAZyme). Core, shared genome (6-48 strains), and unique genome (<5 strains) illustrates distribution analysis of functions across all *Candida* species. g, Analysis of enzyme function of glycoside hydrolase family across all *Candida* strains. h, Breakdown of cell wall composition of core *Candida* strains with identification of 49 CAZymes.

The collection of *Candida* strains used to integrate functional annotations can be categorised into commonly invasive, non-invasive (requiring a secondary factor to cause infection, such as co-morbidity, immunodeficiency) based on literature (Fig. 1d, Supplementary Table 5, Method). These representative samples of *Candida* were integrated into the functional analysis framework, with a total of 49 *Candida* species annotated with BioFung, Protein families (Pfam)^63^ and Carbohydrate-Active enZyme (CAZyme)^64^ databases. We applied BioFung using the UCLUST algorithm, to establish core genome features (found in all *Candida* species) and accessory genome features (shared or unique functions) ^65^. Intra-strain analysis of *C. albicans* across 24 strains sequenced showed largely conserved metabolic pathways and CAZymes (Extended Figure i-j). In covering KEGG metabolic orthologs, clustering analysis determined a larger number of accessory features of metabolism compared to core characteristics seen in all *Candida* strains (Fig. 1d, Extended Fig. 1f-h, Method).

### Identification of global functional annotation profiles in *Candida*

We next determined the CAZyme profile by mapping the 49 *Candida* protein sequences to the dbcan2 database^64^. Doing this allowed us to infer molecular enzyme function^66^. CAZymes are vital enzymes involved in the metabolism of complex carbohydrates. Approximately 205 unique CAZymes were identified in all *Candid*a strains, with various functions (Fig. 1f, Supplementary Table 6, Method). From core *Candida* genome analysis, annotated enzymes were distributed across 6 active families: providing an assortment of enzymatic functions. The glycoside hydrolase (GH) family showed the highest degree of core coverage (Extended Fig. 2a), with much of the GH family activity in starch and other storage carbohydrates substrate-converters (Fig. 1g).

**Figure 2.**
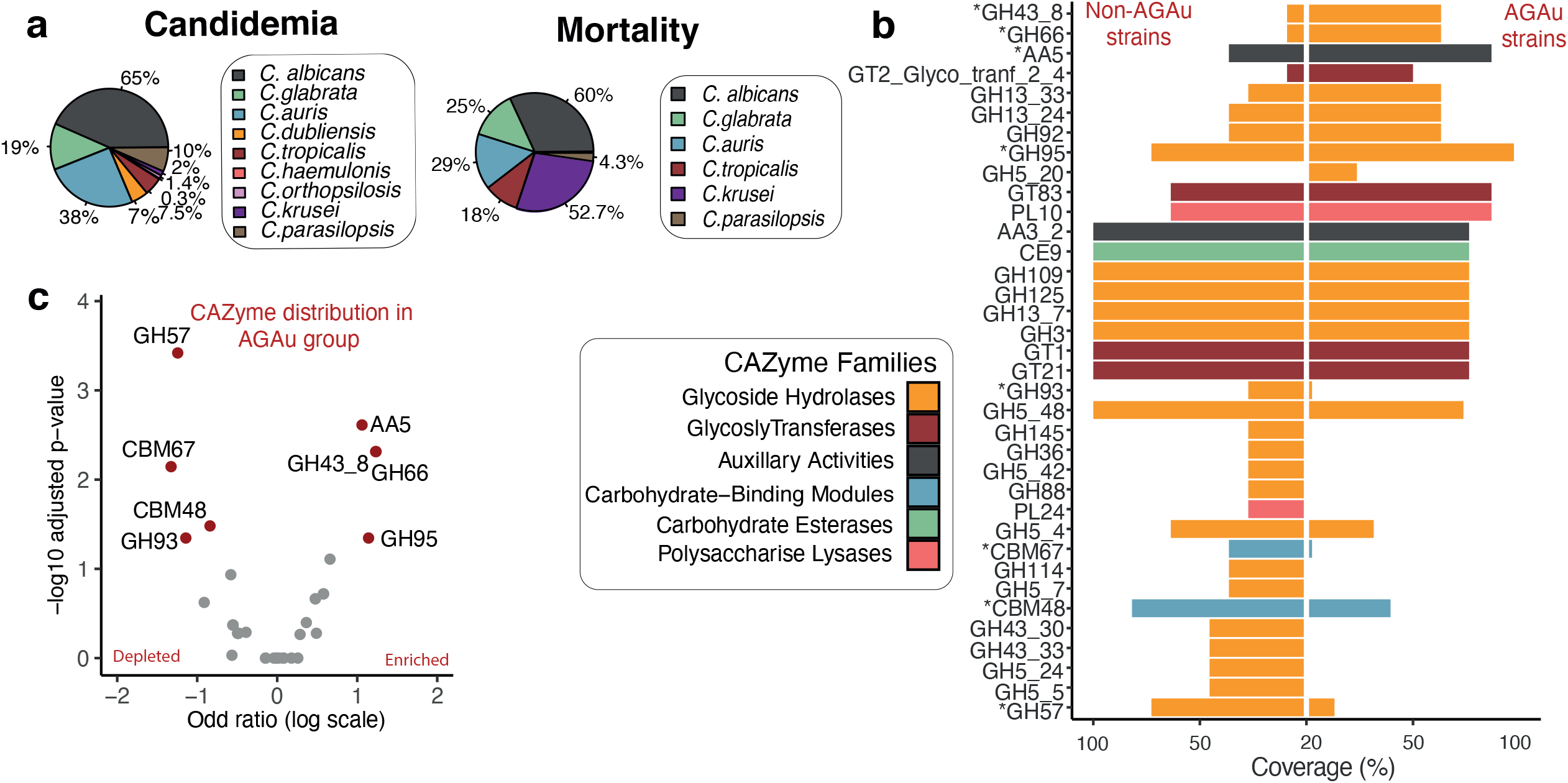
a, Contribution of individual *Candida* species to candidemia and mortality. The impact of each species in AGAu species’ grouping is attributed in this study (literature-based) ^16–20^. b, CAZyme analysis across AGAu and non-AGAu groups. Asterisk connotes statistically significant CAZyme. c, CAZyme enrichment and depletion in AGAu strains. Volcano plot showing statistically significant evidence of a relationship between AGAu strains (chi-squared<0.05, in red). Odds ratio applied to distinguish the enriched and depleted effect in invasive strains.

Additionally, xylan and sugar carbohydrate utilisation were the dominant functions in the accessory genome (Supplementary Table 6). Cell wall composition is a crucial virulence factor and assessing CAZyme components of cell wall substrate converters has been extensively researched^67^. Here, we reveal the presence of pectin lyases, glycan lyases, chitin lyases and mannan lyases (Fig. 1h). Pectin substrate conversion enzyme has been identified as the core feature of *Candidas’* functional cell wall enzyme, though previously only reported in *Candida bodinii*^68^and frequently seen in the fungal plant pathogen, including *Aspergillus Pencillium*^69^. Alongside ß-glucan, mannan and chitin carbohydrate enzyme profiles, *Candida* cell wall activity includes pectin enzyme activity (Supplementary Table 7).

In addition, we identified 1,182 Pfam clans from all *Candida* strains and re-categorised them into 14 functional clans (Supplementary Table 8). Pfam domain annotation indicating genetic information processing, cell machinery, and metabolism was among the most extensive Pfam domains exhibited (Extended fig. 2b). We assessed the diverse functional association of protein domains by analysing core functional clans, and determined similar patterns of dominance for carbohydrate, amino acid and lipid processing-associated domains (Extended Fig. 2c).

### The functional and metabolomic activity of clinical AGAu *Candida* strains

Next, to better explore and understand the link to metabolism and pathogenesis, we clustered the groups based on invasive nature of particular species, linked to contributing to a high percentage of mortality and candidemia (Fig. 2a). *C. <u>a</u>lbicans, C. <u>g</u>labrata* and the emerging invasive species *C. <u>au</u>ris* were grouped together (AGAu cluster). Alternative *Candida* species termed non-AGAu group include opportunistic species that require virulence factors or a defective immune system to cause disease pathology as well as environmental *Candida* species. The AGAu cluster contains those *Candida* species most commonly associated with clinical pathology, contributing to a higher percentage of mortality and candidemia ^16–18,53– 55,70^. This classification of AGAu is analysed and discussed throughout the rest of this paper.

We compared the CAZyme profile coverage of AGAu and non-AGAu groups (Fig. 2b-c, Supplementary Table S9). The CAZyme GH43_8 (substrate-conversion of α-L-arabinofuranosidase/β-xylosidase ^71^) was significantly enriched in AGAu possibly involved in the breakdown of complex glucans (Wilcoxon signed-rank test, P-value <0.05, Method)^72^. The identification of significant CAZymes in the AGAu cluster showed carbohydrate conversion of xylan (GH43_8), mucin (GH95), cellulose (GH66) and copper oxidase family (AA5) ^73–75^. Interestingly, GH66 associated with human oral plaque formation^73^ and AA5 enzyme has been reportedly linked to fungal defence ^76^. CAZymes seen in non-AGAu species showed carbohydrate-binding module families involved in sugar, polysaccharide and cell wall breakdown, including CBM48.

The AGAu species are morphologically diverse, (*C. albicans* is dimorphic, whilst *C. glabrata* and *C. auris* are not), they are all potential pathogens. Therefore, we assessed metabolic pathway enrichment in KO annotations of both AGAu and non-AGAu *Candida* strains (hypergeometric test and Wilcoxon signed-rank test, p-value < 0.05, Method). We found functional evidence of pathways present in both clusters, indicating a genetic potential for all *Candida* strains to undertake similar metabolic trajectories, as no differences were observed between cluster group in pathway analysis (Fig. 3a). All *Candida* strains notably revealed encoded pathways facilitating carbohydrate catabolism within the system; thus, potential to drive increasing metabolic activity, for example through fructose and mannose metabolism. We identified significant enrichment of amino acid metabolism, including arginine, proline, cysteine and methionine metabolism. We also observed significant levels of fatty acid biosynthesis and glutathione metabolism, which have previously been associated with virulence mechanisms^77–80^.

**Figure 3.**
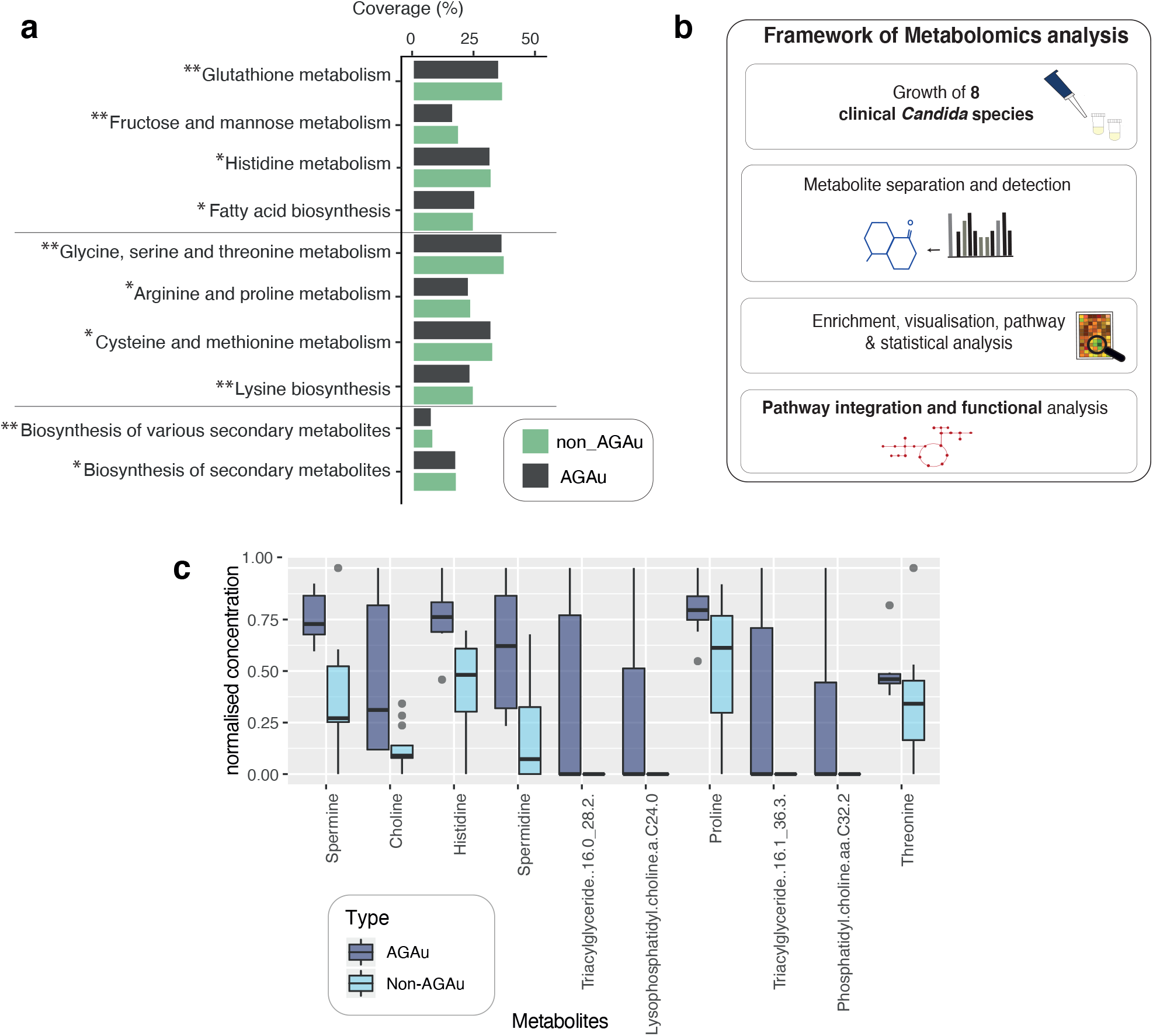
a, Pathway enrichment in 49 *Candida* strains. Coverage of pathways in AGAu and non-AGAu groups indicating genetic pathway potential (hypergeometric testing and Wilcoxon signed-rank test <0.05). b, Framework outline of metabolomic analysis of *Candida* species’ metabolism levels of bioactive metabolites in *Candida* strains exhausted media. Computer simulations were performed for pathway analysis, and statistical approach was applied for candidate metabolites that have a potential influence on the host. c, Significantly enriched metabolites detected in metabolomics in AGAu and non-AGAu groups (Wilcoxon rank-sum test <0.05).

### Metabolomics revealed key metabolic pathways assimilated by AGAu group

To elucidate the metabolic trajectory taking place by each cluster group, we performed metabolomics on a collection of 7 clinical *Candida* isolates, representing the diverse pathogenic species (Fig. 3b, Method). These clinical samples were previously isolated from patients infections ^53,81–84^ and were used to evaluate *in vitro* the critical metabolic activity predicted by our functional analyses (Extended fig. 3a). These *Candida* isolates were representative of both AGAu and non-AGAu grouped species. The metabolomics data were used for partial least squares discriminant analysis (PLS-DA), which resulted in distinct cluster separation of significant analyte classes (Extended Fig. 3b-d, Method). The PLS-DA model identified distinct high metabolites features uniquely in AGAu *Candida* species (Extended Fig. 3e). For example, histidine metabolite production in AGAu species supports this metabolite role in systemic infection and is potential an anti-fungal target^85,86^. We also identified choline-derived metabolites (choline, phosphatidyl-choline and lysophosphatidylcholine) as increased in certain AGAu species (Fig. 3c). We identified phosphatidylcholines analyte class as contributing the highest number of features across selected clinical *Candida* species (Extended Fig. 3d), which has been observed previously in the hypervirulent *C. albicans* (SC5314) strain^38–41^. Lastly, spermine and spermidine, were found to be significantly associated with AGAu strains, indicating polyamine metabolism could play a functional role in the increased association with disease pathology of these strains.

### Integrative global metabolic map of of AGAu *Candida* species

Having identified these pathways in silico, we next determined gene expression levels of essential polyamine (SPE11, SPE3), choline (CKI1, TAZ1) and fatty acid (CEM1) pathways in *C. albicans* from the AGAu cluster to validate these findings (Extended fig. 4d, Supplementary Table 10 and Methods). All 5 genes showed expression substantiating the activity of these pathways. Our conclusions of key pathway associations draws importance of nitrogen sources, specifically the metabolism of amino acids, in this process (Fig. 4), although to date, *Candida* species pathogenesis is better known to be driven by carbon sources^22,37,89,90^. AGAu species exhibited increased levels of metabolites in the choline pathway, polyamine and fatty acid biosynthesis pathways that are primarily propagated through arginine, cysteine and methionine pathways (Extended Fig. 4a-c). Based on integration of computational and experimental data revealed fundamental metabolic pathways applied AGAu species providing a link to the major advantage shown by AGAu species across the human body with considerable contributions to pathogenesis. These important pathways include polyamine, choline and fatty acid biosynthesis. For instance, the polyamine pathway is thought to be involved in *Candida* cell proliferation, and in turn, causes host cellular dysfunction by modulating acetylation levels of aminopropyl groups and inducing autophagy, thus increasing cell life span^89,91,92^. We observed fatty acid biosynthesis production with large numbers of metabolites of triglycerides featuring in PLSDA and family have previously been reported to promote germination and virulence of AGAu *Candida* strains (Extended Fig. 3d)^93,94^. Further, fatty acid biosynthesis is vital in fungal cell membrane viability, energy storage, signalling, and cell proliferation – all functions critical in pathogenesis^95–98^.

**Figure 4.**
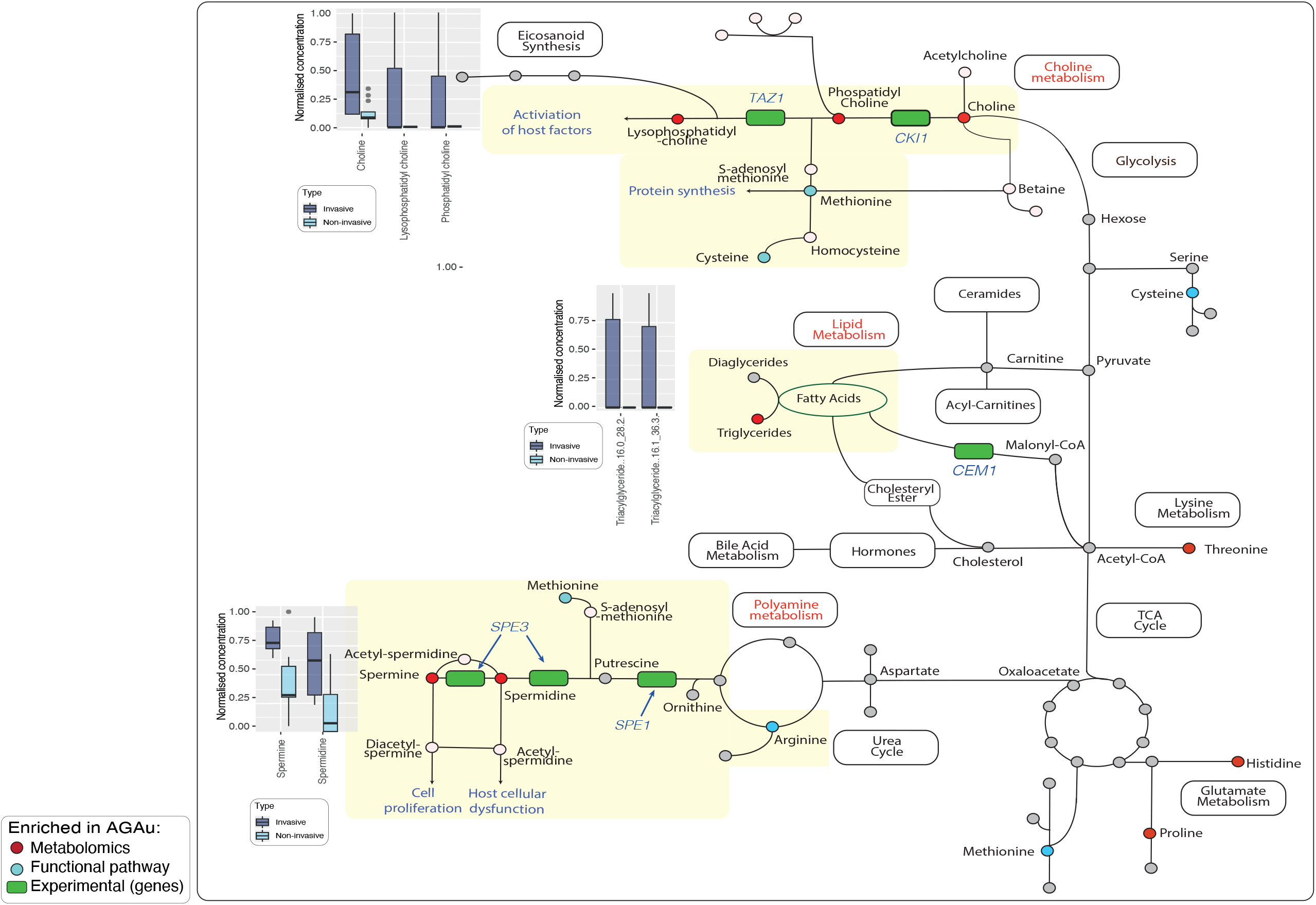
Global map metabolism in AGAu *Candida* strains. Enriched pathways observed in functional annotation and further validated by metabolomics with key metabolites markedly significant (Colour in pathway red identify significant metabolites, blue indicate significant pathway enrichment).

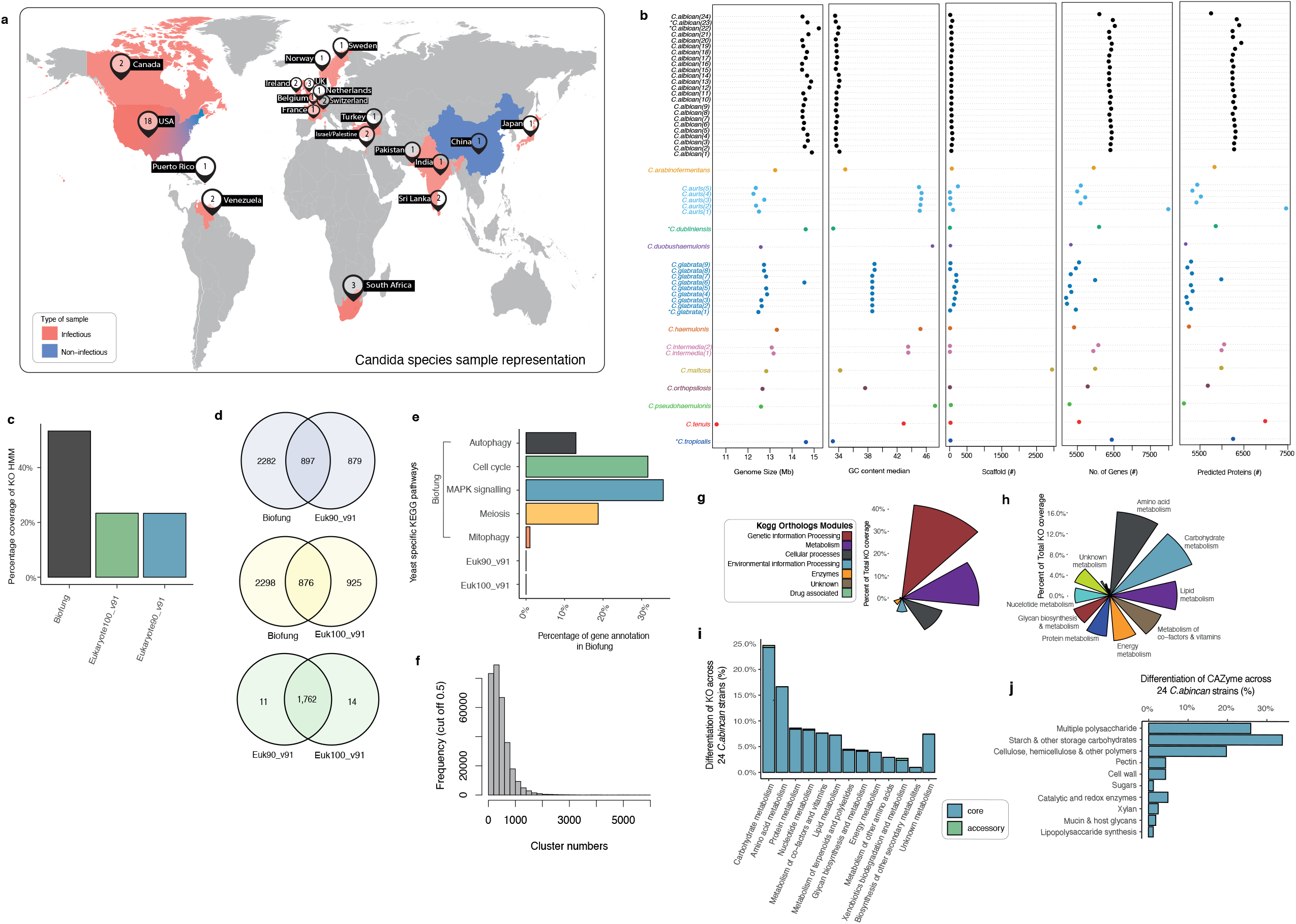

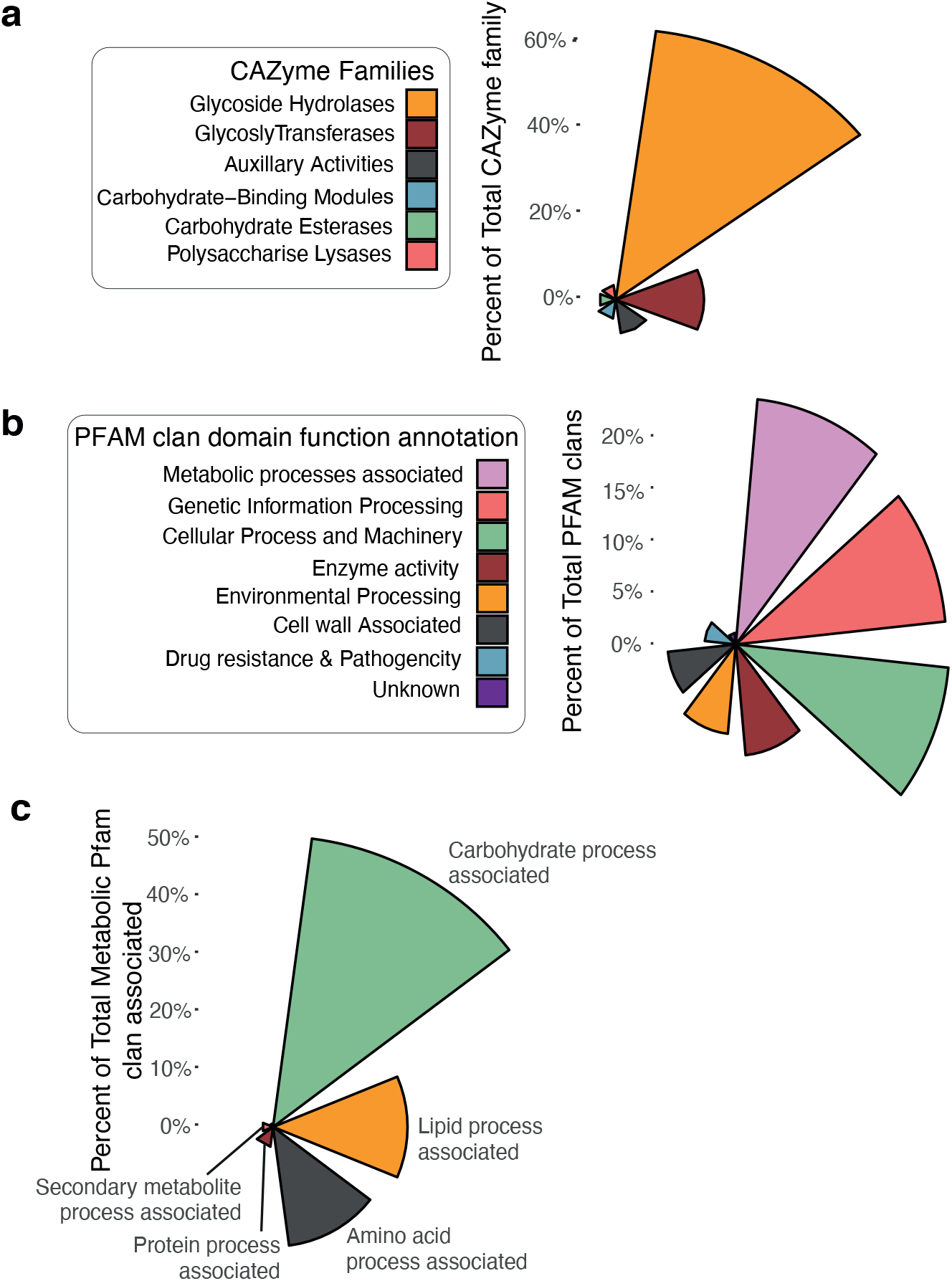

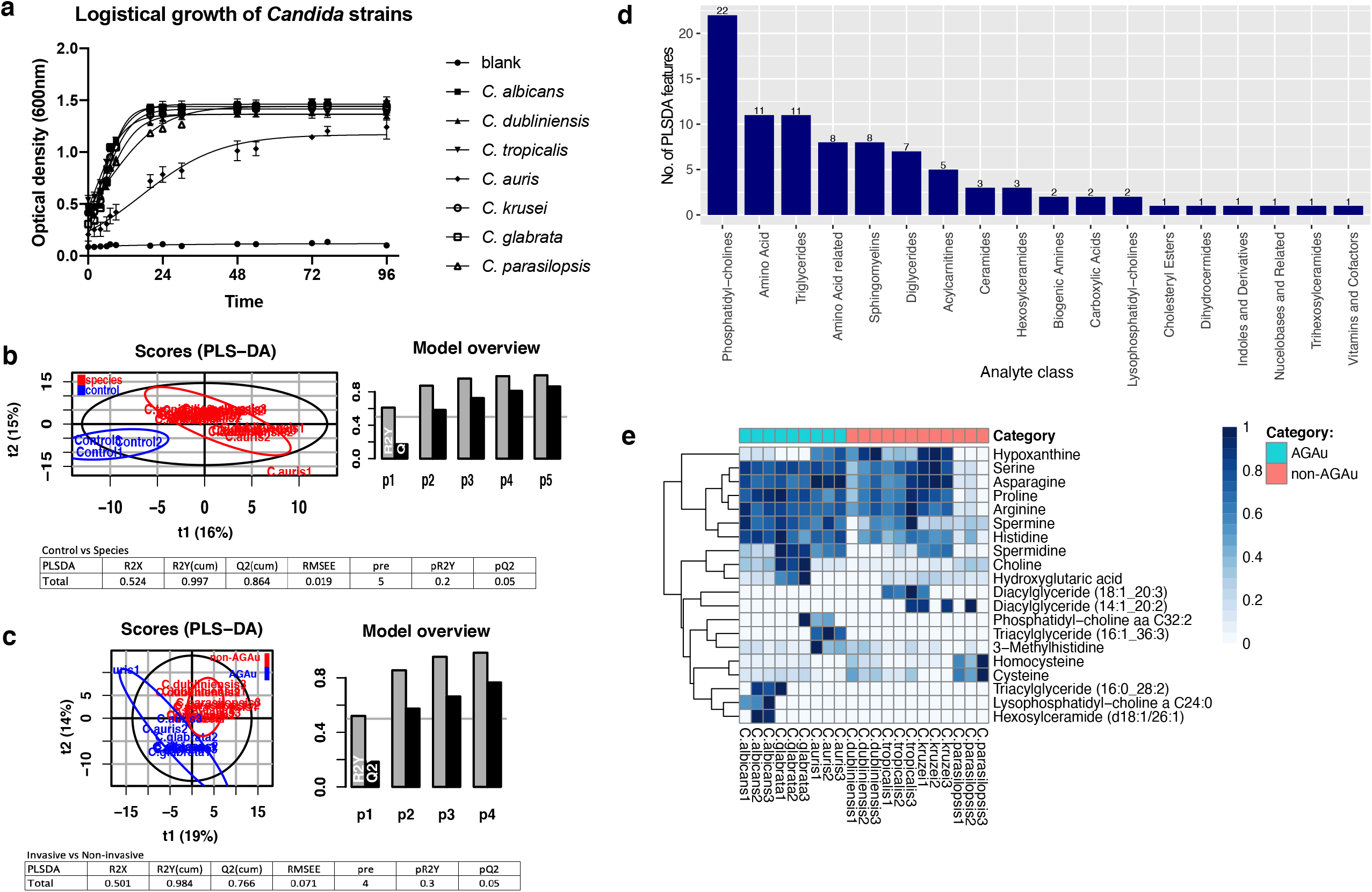

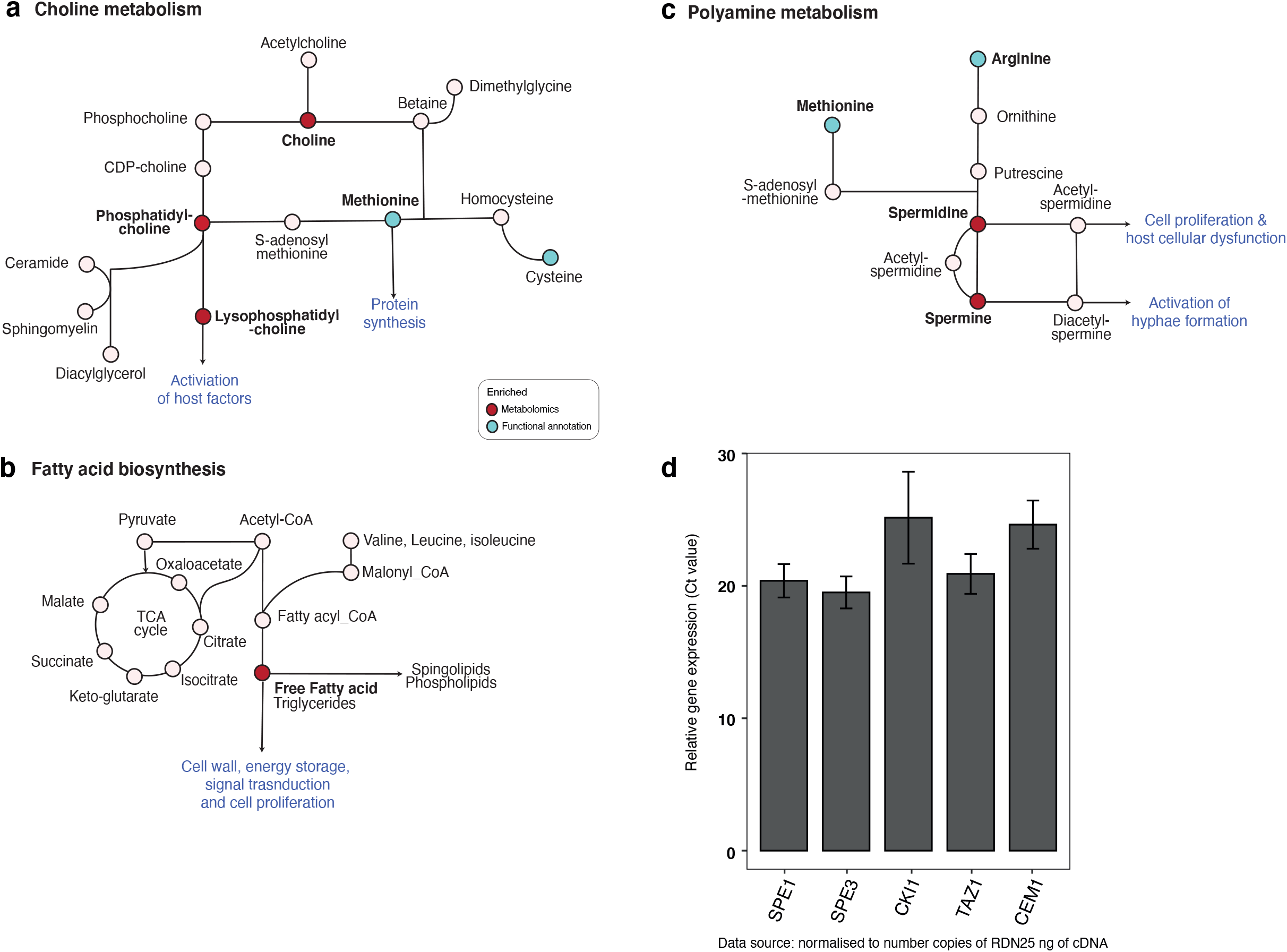

## Discussion

BioFung was used for building metabolic maps of key *Candida* strains and the database provides the mycology community with a resource allowing them to dive more deeply into all fungal species’ metabolic capability based on protein encoding genes. Development of data generation technologies development, tools and database for fungal species is currently in its infancy, despite significant advances in these areas for bacteria and archaea. This database enables detailed mechanistic annotations to optimise our understanding of fungal species. to do this, it uses HMM to provide high specificity for fungal annotation. To do this, it is currently the best database available for KEGG-based functional annotation of fungi (Extended fig. 1b-d and Supplementary Table 2-4).

Analysis of annotations allowed us to identify the influential AGAu group of *Candida* strains, highlighting critical metabolic pathways in these strains. In doing so, we developed increased understanding of the metabolism of these strains through integrating multi-omics and experimental data. BioFung can be used extensively to better understand individual fungal species’ metabolic pathways but can be extended to explore metabolic interactions between fungi, other organisms, and within the host-mycobiome.

Using this tool in combination with metabolomics validation, the AGAu *Candida* strains appears to be employing specific activity of amino acid metabolism. This shows a degree of metabolic plasticity indicative of fungi, where secretion of these metabolites associated with these pathways aiding in better adaptability to growth, virulence factor production, hyphae and biofilm formation^39,99^, enabling more effective adaptation to a wide variety of environments and habitats. Amino acid metabolism has been proposed as an alternative energy source in stress responses and an alternative to carbon sources for growth. We demonstrate here that all Candida species have an amino acid pathways to employ metabolic remodelling (Fig. 3a)^39,42,89^. However, integrating metabolomics from strains grown with abundant nutrient source, shows that the AGAu group as opposed to other species retains significantly more active in the production of polyamine and choline metabolites, which requires the use of amino acid metabolism for production (Fig. 4). The AGAu group employs arginine, methionine and cysteine metabolism and more extensive exploration and experimental data needed to understand the causal effect of amino acid metabolism. For instance, we have identified a confirmed target pathway for anti-fungal drugs with glutathione metabolism (GSH), attributed to fungal mitochondrial maintenance, preservation of membrane integrity, regulation of transcription factors in stress response and protection against reactive oxygen species. Reducing activity of GSH is under investigation as supplementary aid for anti-fungal drugs (azoles and echinocandin) against resistant strains ^100–103^. This finding verifies that pathway enrichment analysis echoes feasibility in clinical relevance within the host.

Among these metabolites identified are the polyamines, including spermine and spermidine. Polyamines play critical roles in normal cell physiology. Spermine is essential for *Candida* hyphal formation, playing a pivotal role in *Candida* invasion^104^. Spermidine, meanwhile, drives genetic modification in fungi by regulating cell cycle and translating the modification of eukaryotic initiation factor (eIF)^105–107^. Excessive polyamines prolong yeast survival via delayed DNA degradation, increasing the likelihood of mutations that could contribute to the development of anti-fungal resistance ^108,109^. These mutations are an important consideration, given that *C. glabrata* and *C. auris* are heavily associated with rising anti-fungal resistance^110–112^. Polyamines have also been shown to be anti-inflammatory, depending on the microenvironment, potentially explaining the additional benefits of secondary metabolites to *Candida* species by modulating host immune responses^113^. The use of polyamines is not limited to fungi. Bacteria use polyamines to create and maintain biofilms in order to withstand host defences as well as promoting cancers^114–116^. Viruses use polyamines to promote cell proliferation, thereby promoting their propagation and spread. Intervention in polyamine synthesis has a high degree of potential as a target for antimicrobials. DNA viruses upregulate polyamine synthesis in host during infection and blocking polyamine synthesis is a strategy used in broad-spectrum anti-viral^117,118,119^. These examples along with our findings here indicate that manipulating polyamine secretion from *Candida* species is a realistic target for therapeutic intervention of associated diseases.

Choline metabolism is a critical function for both microbial and host physiology, as demonstrated by the increase seen in AGAu Candida species’ related metabolites. Disruption of phospholipid biosynthesis in fungi can occur through inhibition of phosphatidylcholine synthesis, showing preventing virulence within the systemic mice model^87,88,120,121^. Further, acetylcholine is essential in the formation of the chitin wall characteristic of fungi^122,123^. Along with the bacteriome, *Candida* species contribute towards host acquisition of choline. As understanding of choline metabolism is in its infancy, further investigation of host-mycobiome interactions is needed, potentially providing insights for repurposing potential therapeutic intervention. For instance, lack of choline in humans drives liver dysfunction due to the accumulation of lipids within hepatocytes, which can lead to fatty liver diseases and even hepatic liver cancer^124–128^.

Functional analysis indicates that both the AGAu and non-AGAu groups show a great degree of metabolic plasticity (Fig. 3a)^56,59,60,129^. The only functional difference seen between groups was with CAZymes. The AGAu group presented with GH66, which has previously only been associated with the human oral microbiome and as a potential marker for plaque formation^73^. Given that *C. albicans* is constituent of oral plaque, this is constituent with clinical data (Fig. 2b-c) ^130^. The finding of GH43_8 remains relatively inconclusive in function but was recently detected in bacteria as β-galactofuranosidase^131^. Although the modes of action for both GH43_8 and AA5 are currently unknown their enrichment in the AGAu group may provide a function-targeted biomarker for *Candida* infection^132,133^. We also highlight fatty acid biosynthesis pathway in AGAu species with significant levels of triglycerides’ production detected (Fig. 3c). Fatty acid synthesis has been identified in *Candida* species previously, with focus on OLE1, FASI & FASII genes as key indicators to pathogenesis and virulence^77–80^. This validates the notion of targeting fatty acid biosynthesis pathway within *Candida* species to disrupt *Candida* overgrowth in the host.

Our study has addressed the need for functional data and tools for fungal species by developing the BioFung resource using the KEGG database. This enables detailed mechanistic pathway analysis of fungi. Our integrative analysis of the AGAu group (associated with the disease pathogenesis) highlighted key pathways that potentially increase virulence and have associating effects in the host. We hypothesise that these markers can aid in identifying routes for intervention in invasive infection. We suggest polyamine, choline and fatty acid biosynthesis metabolism as inception for further investigation. The presence of these metabolites from AGAu *Candida* species potentially directly affects host homeostasis within the mycobiome and adverse effects on humans during infection. As such, the AGAu *Candida* species’ metabolic reprogramming may present a novel method of controlling interaction and infection with these fungi. Finally, we focus on fungal metabolism exploration and distinctively towards amino acid metabolism, playing a more significant role in virulence and pathogenicity.

## Code availability

BioFung is public open access database that can be downloaded at: https://www.microbiomeatlas.org/downloads.php. The instruction and the pipeline scripts for BioFung can be found at our GitHub repository https://github.com/sysbiomelab/BioFung.

## Acknowledgment

This study was supported by Engineering and Physical Sciences Research Council (EPSRC), EP/S001301/1, Biotechnology Biological Sciences Research Council (BBSRC) BB/S016899/1, Science for Life Laboratory, Swedish National Infrastructure for Computing at SNIC through Uppsala Multidisciplinary Centre for Advanced Computational Science (UPPMAX) under projects SNIC 2020-5-222, SNIC 2019/3-226, SNIC 2020/6-153 and King’s College London computational infrastructure facility, Rosalind (https://rosalind.kcl.ac.uk) for high performance computing. We thank Professor Bernhard Hube for kindly sending *C. parapsilosis* strain.

## Author contributions

N.B, D.M. and S.S. conceived the project. N.B. performed sample preparation, metabolomics, gene expression data preparation and extraction protocols for the paper. N.B. developed the pipeline, analysis and made the figures. S.L. advised on design, statistical and functional analysis of the data. A.P.and S.S.N. processed qPCR platform. N.B. wrote and drafted the manuscript. J.N. and M.U. provided critical feedback on the data and manuscript. All authors read, edited and reviewed the manuscript.

## Conflict of Interest

The authors declare no competing financial interests.

## Methods and Materials

### Genome sequence collection

Genome sequences of 49 *Candida* species were collected from NCBI database with version release 45 of Ensemble Fungi (date accessed: April 2019)^134,135^. Supplementary information of sample strain, genome ID, ENA ID, Biosample ID, sequence platform, year of collection, sample location of collection, sample tuple and available biological annotation has been provided in Supplementary Table 1. The quality of the sequence was checked to look at GC content, scaffold and genome size (see Supplementary Figure 1b). Nucleotide sequences used with ANI to determine the phylogenetic relationship and determine differences between strains using Pyani package^136^.

### Construction of fungi-specific functional database (BioFung)

Kyoto Encyclopedia of Genes and Genomes (KEGG) database were downloaded for the investigation of all 128 fungal species (3GB file size) from eukaryote database (5GB file size) (downloaded on August 2019)^137^. Around 1,210,746 genes, which are annotated with 4,717 KEGG orthology (KO), were selected among 128 fungal species genes. There were 6071 fungal genes missing sequence to place into KO, and 105 KO failed in multi-sequence alignment due to default settings (minimum of 3 genes sequence required). Those genes per each KO were performed multiple-sequence alignment by ClustalW and generated Hidden Markov Model (HMM) profiles using the hmm-build function of HMMER software (Figure 1A for workflow and supplementary 2a for coverage)^138,139^. Our database of fungal-specific HMM models of 4,722 KOs was freely shared via Github repository (https://github.com/sysbiomelab/BioFung). Missing KO from fungal species was not added due to missing gene sequences from KEGG, or the low number of sequences per KO (<3), thereby failed to perform ClustalW alignment (See details in Supplementary Table 4). Quality check was performed by comparing BioFung HMMs to pre-trained HMMS for eukaryotes (euk90 and euk100 version 91.0) from Raven Toolbox ^57,58,^ and we observed that BioFung coverage was much higher than both eukaryote profiles (Supplementary figure1c).

### Functional annotations of individual fungi species

Fungal KO annotation of each species was performed by HMM scanning of BioFung HMM models by HMMER software. An in-depth exploratory analysis was performed by manually checking KO annotations of individual species deep. Pathway abundance for AGAu and non-AGAu species was performed using KEGG pathway annotations. Hypergeometric testing of pathways with significance confirmed using Wilcoxon signed-rank test (<0.05). CAZymes annotations were performed by mapping *Candida* protein sequences using HMMs of dbCAN2 database^66^. Substrate conversion of CAZyme families was checked based on literature review^69,140–146^.

*Candida* protein sequences to map against Pfam-A families using HMMs, that are fully annotated and curated above a threshold^63^. Pfam clans’ annotations were sub-set into a broader annotation based on a reported standard function of protein domains (please see Supplementary Table 8).

### Contrasted functional annotation of *Candida* species

Presence and absence of microbial annotations, i.e., prevalence, was tested for significance based on condition using Chi-squared tests and odd ratio. Percentage coverage of each was also tested between AGAu and non-AGAu *Candida* species. Contrasted functional annotations were checked on individual strains and placed into presence/absence to perform chi-squared for significance (<0.05), and the odds ratio was performed to identify enriched and depleted in AGAu samples. Additional significant functional annotations are seen in AGAu cluster (Supplementary Table 9).

### Clustering of protein sequences

Core, shared, and unique proteins were identified based on sequence similarity by a clustering approach called UCLUST algorithm^65^. In short, UCLUST algorithm was applied to identify similarity in protein-encoding gene sequences by clustering and unique protein sequences were identified if included in singleton protein clusters. Core proteins were identified if corresponding proteins from all 49 species were included in the same cluster. Shared proteins were selected if they did not belong to unique and core proteins. Protein sequence clusters were selected based on a threshold 0.5 for representative seed sequence, a default threshold in UCLUST software (Extended fig. 1h). Based on definitions of the core, shared, and unique proteins, we were able to determine the core, shared and unique annotations for KO and CAZymes, accordingly.

### Strain growth

8 strains of *Candida* species (*C. albicans (*SC5314), *C. dubliniensis* (CD36), *C. tropicalis* (CBS94), *C. glabrata* (CBS 138), *C. auris* (47477), *C. parapsilosis* (73/037), and *C. krusei* (CBS573). Strains were grown in liquid sabouraud dextrose broth (Thermo Scientific-Oxoid microbiology, UK)^147^. All strains were grown in 50ml falcon tube in a shaking incubator 95rpm at the temperature of 25°C to encompass all growth rates. Timepoint measurement of growth was taken to measure the exponential and stationary phase of the optical density of 1 at 600nm absorbance (iEMS Ascent absorbance 96-well plate reader).

### Collection and targeted metabolomics on fungal extracellular matrix

Mid-exponential phase indicates bioactive metabolites and time points for the extraction of extracellular metabolites (see Supplementary Figure 1a). 500µl of extracellular medium, proximity to the pellet was removed from growing fungal cells. Samples were placed through 20µm Whatman filter and snap-frozen in liquid nitrogen. Targeted metabolomics performed using the MxP Quant500 kit (Biocrates, Austria). Partial Least Square – Discriminant Analysis (PLS-DA) was performed on targeted metabolomics of fungal extracellular matrices and media as control, using *ropls* package^148^. First, PLS-DA was performed to distinguish between *Candida* samples and control (media). Further, PLS-DA was performed to distinguish between AGAu species and non-AGAu *Candida* samples. PLS-DA indicated a significant difference between AGAu clusters. Further analysis of metabolite concentrations of targeted metabolomics was normalised, and the Wilcoxon rank-sum test was performed to identify critical metabolites and pathways (<0.05).

### Validation experiment

RNA was extracted from 3 biological repeats *C. albicans* (SC5139) using RNA Qiagen Powersoil kit adapted with bead beating with interval placement on dry ice and additional 100µl of isopropanol. DNAse cleanup performed using RNA clean-up and concentration kit (NORGEN, Biotek corporation). Primers designed for specific amplification of genes SPE1 targeting Ornithine Carboxylase, SPE3 gene for spermidine synthase, CKI1 specific for bifunctional choline kinase/ethanolamine kinase, TAZ1 gene focused on lysophosphatidylcholine acyltransferase) and CEM1 gene target for fatty acid synthase (Supplementary Table 10 for primer information). These primers are specific for *C. albicans. O*ther *Candida* species only predicted gene ontology-based on *C. albicans* and *Saccharomyces* annotation (http://www.candidagenome.org/cgi-bin/GO/goAnnotation.pl?dbid=CAL0000224407&seq_source=C.%20auris%20B8441). Conventional RT-qPCR was performed to identify the expression of these critical pathways for samples, two standard curve analysis with RDN25 which encodes the 25s rRNA subunit.

## Figure legends

**Extended Figure 1**. a, Map of *Candida* species collection. Indication of global representation of samples. b, Sequence quality assessment. Genome size variation, GC content, number of contig and outcome of quality of the sequence. c, Comparison of BioFung to available euk90 and euk100 KEGG profiles. Euk90_v91 and euk100_v91 are versions of pre-trained hmm for eukaryote database^62^. Coverage of BioFung was higher than of available hmm. d, Venn diagram displays the overlap of annotation. The overlap between BioFung vs euk90_v91, BioFung vs euk100_v91 and comparison of both eukaryote profiles from all candida gene annotation. BioFung still has an enormous scope of unique KO, and small differences between eukaryote profiles can be observed. e, Validation assessing coverage of the yeast-specific pathway. We performed pathway analysis of KO annotations containing yeast-specific cell pathways. Observation of no yeast specific pathways was found in eukaryote annotation. f, Core percentile coverage of functional annotation. First instance, distribution of KEGG module categories reflect functions seen in most microbiota species. Illustrates the extensive functional capability of *Candida* within each species. g, Core KEGG metabolism exploration in *Candida* species. Amino acid, carbohydrate and lipid metabolism contributes the greatest make up of gene function in *Candida* species. This indicates the contribution of different metabolic potential from *Candida*. h, Uclust quality assessment by looking at the frequency of cluster numbers at 0.5 cut-offs. The cut-ff was inclusive of signature clusters capture and a broad range of core clusters. i, Intra-strain metabolism analysis of *C. albicans* indicating genome differentiation. j, Clustering of 24 *C. albicans* strains of Carbohydrate Active enzymes demonstrating indistinguishable changes in genome.

**Extended Figure 2** a, Coverage of CAZyme indicates glycoside hydrolase family is the largest contributor in all *Candida* species. b, Pfam coverage indicates that metabolic processes, genetic information processing and cellular process & machinery provide the immense repertoire of protein function in all *Candida* species. c, Core Pfam metabolic process analysis in *Candida* species. Carbohydrate, amino acid, and lipid-associated processes are the largest metabolomic protein function in *Candida* strains.

**Extended Figure 3** a, Growth curve of strains grown in the laboratory. All the strains were grown, and timepoint measurement of OD1 at 600nm was taken to obtain the mid-exponential phase. b, Targeted metabolomics with PLS-DA analysis differentiating from media. PLS-DA score indicated a precise fitting performance of clustering media distinctly from *Candida* strains. c, Targeted metabolomics PLS-DA analysis based on AGAu and non-AGAu species. PLS-DA score plot indicated a fitting and predictive performance (2 latent variables, *R*^*2*^*X* = 0.501, *R*^*2*^*Y* = 0.984, *Q*^*2*^ = 0.76). d, Analyte Class component features in PLS-DA analysis. A significant number of differential components between invasive and non-invasive strains. e, Heatmap of significant metabolites. Identified from PLS-DA, top 20 metabolites differentiated in AGAu and non-AGAu based on Wilcoxon rank-sum test (>0.05).

**Extended Figure 4** a, Choline metabolic pathway. Concise choline metabolism and its affect. b, Fatty acid metabolism. Concise fatty acid biosynthesis metabolism and its affect. c, Polyamine metabolism. Concise polyamine metabolism and its affect. d, Quantitative RT-qPCR validation of pathways with *C. albicans*. Indicating pathway genes (SPE1, SPE3, CK1, TAZ1, CEM1) compared to their relative expression in *Candida albicans*, representative of AGAu species cluster. (n = 3 biological replicates. Display the relative RNA expression level (Ct value)).

**Supplementary Table** 1: Information on database collection of 49 *Candida* strains. In-depth data collection of strain retrieval from a database, including strain id, genome id, sample type, origin country, year of collection, annotation availability.

**Supplementary Table** 2: Fungal species in KEGG database. List of 128 fungal genes in KEGG database, species abbreviation, T number, protein-encoding genes and genes converted to KO.

**Supplementary Table 3**: The building of BioFung. List of fungal-specific KO available in KEGG database, genes available per KO, missing genes from KO, missing sequences of genes unable to include and failure of multi-sequence alignment in creating BioFung.

**Supplementary Table 4**, Missing information from the KEGG database. List of genes in fungal species had missing sequences and was unable to collate in profile.

**Supplementary Table 5**, AGAu strain categorisation. Based on sample names, grouping, location of sample extraction and reference for strain.

**Supplementary Table 6**, The categorisation of Pfam clans. Categories were based on clan function to the associated processes; note clans can fit into many functional categories.

**Supplementary Table 7**, Significant of functional annotation in enriched or depleted in AGAu group. Some annotations are statistically significant with the odds ratio of falling within the scale of 0 and 1. They have not been included in the figure but remain relevant. These have been captured in this table for CAZyme, KO and Pfam.

**Supplementary Table 8**, References for Pectin CAZyme assay confirmation. List of publications confirming the function of pectin. Table 9, References for RT-qPCR validation. List of gene selection, primer sequence and accession numbers for gene expression analysis.

## Notes

### Competing Interest Statement

The authors have declared no competing interest.

